# Alpine peatlands: spatially and temporally complex ecosystems with year-round methane cycling activity

**DOI:** 10.1101/2025.11.07.687222

**Authors:** Sigrid van Grinsven, Sophie Kunz, Florian Jueterbock, Andreas Kappler

## Abstract

Peatlands are well-known emitters of methane. European alpine peatlands share certain characteristics with boreal peatlands, despite being located at temperate latitudes, such as a strong seasonality with snowfall in winter and a short summer and growing season. Unlike boreal peatlands, they experience relatively large temperature fluctuations between day and night and are more likely to be sloping. It is unknown how these factors affect methane dynamics. Furthermore, winter methane dynamics have rarely been studied. We therefore quantified the soil-atmosphere methane flux at an alpine peatland in Austria (1700 m a.s.l), with a focus on the spatial and temporal heterogeneity in this ecosystem. In summer, methane emissions were high (49 mg m^2^ h^-1^), whereas in spring, shortly after snowmelt, both methane uptake and emissions were observed at different locations within the alpine peatland. In winter, a local snow-free patch persisted at the peatland due to the year-round influx of 5°C spring water. We compared the methane flux from this snow-free patch to another alpine peatland which also contained such a snow-free area and observed methane emissions at the one peatland (1.2 mg m^2^ h^-1^) and methane uptake at the other (−0.06 mg m^2^ h^-1^). The input of spring water in combination with the sloping nature of the peatlands resulted in a large spatial heterogeneity, likely as a result of the input of redox-active components such as sulfate by the spring water. The microbial community composition also suggested the presence of active sulfur, iron and methane cycling in the peat soil. Overall, our research shows that alpine peatlands are unique systems due to the year-round spring water throughput, altering biogeochemical cycles and creating local snow-free conditions, with implications for methane cycling.

## 1. Introduction

Peatlands and wetlands are major sources of methane to the atmosphere, due to their anoxic nature and high carbon stocks(Turetsky et al., 2014). Boreal and arctic wetlands have been studied intensively and have been shown to emit large amounts of methane(Kuhn et al., 2025). Although located at a different latitude, boreal wetlands and alpine peatlands share certain characteristics such as a long winter snow cover and a short growing season. It is, however, largely unknown whether alpine peatlands behave similarly to boreal wetlands in terms of methane dynamics.

High latitude (arctic) and high-altitude (alpine) peatlands have predominantly been studied during the summer season due to the practical limitations of visiting these snow-covered places in the winter season. As a result, global models lack inputs on winter methane fluxes from these systems. A recent modelling study pointed to non-growing season wetland methane emissions as the main source of uncertainty in modelling methane emissions(Kuhn et al., 2025).

Temperature, soil moisture content, and plant-effects are all known to play a role in wetland methane emission dynamics(Whalen and Reeburgh, 1996), but their effects are intertwined and complex. Warming experiments with saturated soils have shown that an increase in temperature can result in an increase of both the methane production and methane oxidation potential(Li et al., 2020). The net effect on the emissions depends on the balance between these two. In natural, non-manipulated settings, methane emissions and temperature have been shown to have a direct relationship(Wei et al., 2015). In a recent modelling assessment on the future of methane emissions in boreal wetlands, both temperature and the lengthening of the growing season were important factors causing increased emissions with future climate warming (Kuhn et al., 2025). The vegetation can affect methane emissions by plant characteristics such as root and shoot length and gas transport potential, but the vegetation composition within wetlands itself is also driven by the water availability and hydrologic variables(Harbert and Cooper, 2017). The water table depth in itself can create physical limitations for diffusive gas transport, often leading to anoxic conditions in soils. Several studies have shown a correlation between water table depth and methane fluxes(Chen et al., 2013; Li et al., 2020; Wei et al., 2015), but the mechanisms behind this correlation are often unknown.

We investigate the methane fluxes of two alpine peatlands located in the European Alps (1670 and 2145 m a.s.l.) over different seasons. We also study the high spatial heterogeneity in methane fluxes within these peatlands and use soil and water temperature and chemistry to assess the effect of water transport through the peatlands on summer and winter methane dynamics. Alpine peatlands are unique in their sloping nature, which likely results in short water residence times and high spring water inputs compared to other peatland systems. We show that these features lead to previously undiscovered methane fluxes at naturally snow-free patches in the snow season, and interesting spatial dynamics in the post-snowmelt and summer seasons.

## 2. Material and methods

### 2.1 Site description

The Auenfeld alpine peatland is located at 47°14’27.21”N 10° 7’54.74”E, at 1670 m a.s.l. in Schröcken, Voralberg, Austria. It is ca 0.7 ha in size and located in a wide alpine valley (Fig. 1). The area is part of the Northern Calcareous Alps(GeoSphere Maps). In summer it is used for extensive grazing with cows. In winter, it is located within a well-used ski area but is not part of the ski runs or prepared snow paths. Sampling campaigns at this site took place on 9 July 2024, 20 February 2025, 13 May 2025, 31 July – 1 Aug 2025, and 14-16 August 2025. Unless otherwise specified, gas sampling was always performed between 11:00 and 16:00. The Gleirsch alpine peatland is located at 47° 9’22.43”N 11° 4’40.24”E, at 2145 m a.s.l. in Sankt Sigmund im Sellrain, Tyrol, Austria. It is ca 0.15 ha in size and located in a long and narrow alpine valley (Fig. A1). The bedrock of this area consists predominantly of paragneiss(GeoSphere Maps). In summer it is used for extensive grazing with cows. The sampling campaign at this location took place on 20 January 2025.

**Fig. 1.**
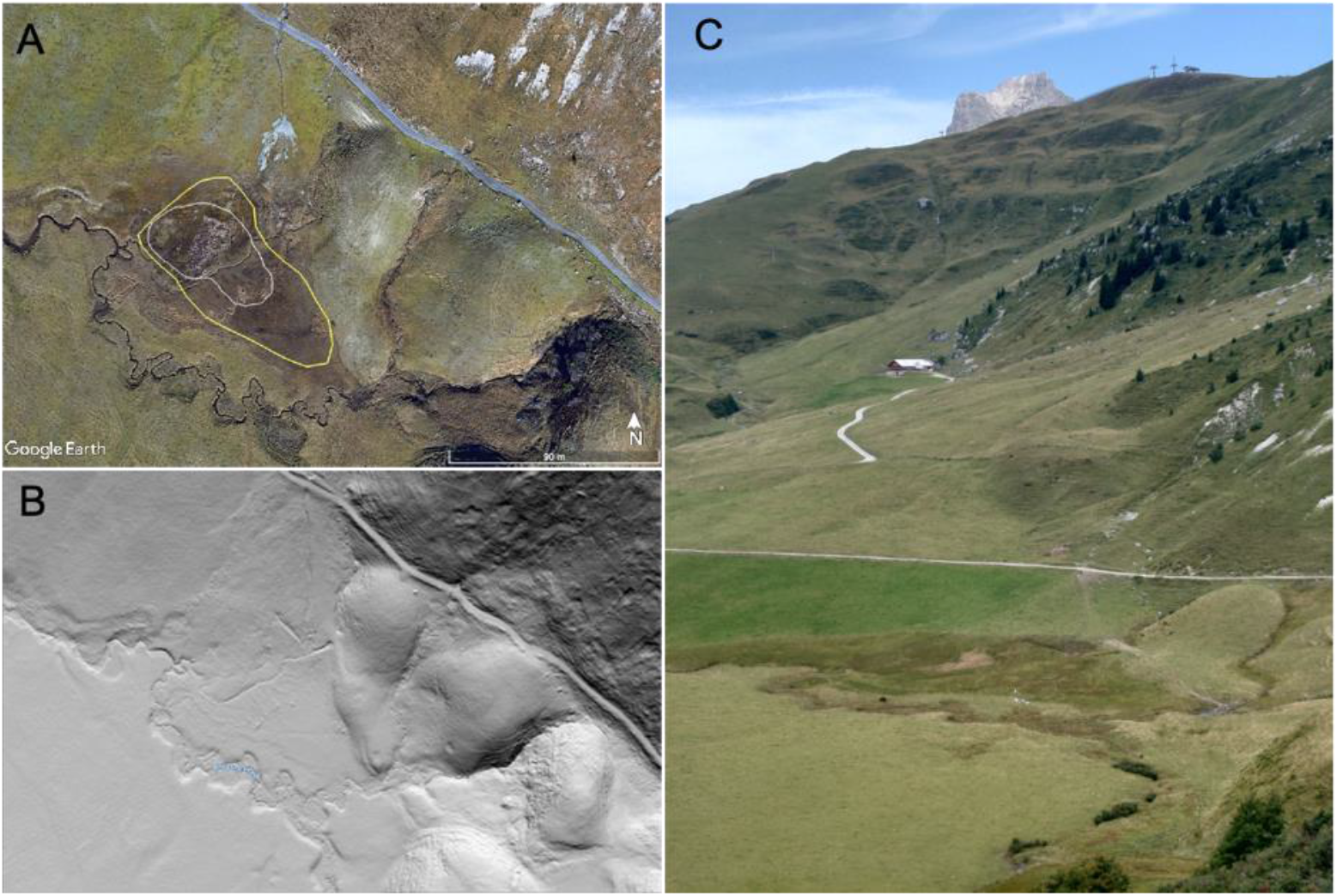
Geographic setting of the Auenfeld alpine peatland site. A) Google Earth imagery with in yellow and white the outlines of respectively the spring/summer and winter sampling areas. B) Hillshade representation of the digital elevation model from the 3rd laser scanning survey of Vorarlberg 2023, 25 cm grid resolution (source: GeoAtlas Voralberg(GeoAtlas Voralberg, n.d.)). C) Picture from August 2025, showing the alpine peatland on the bottom part of the slope, recognizable by its dark green colour.

Maps were created using Google Earth Pro (2025).

### 2.2 Methane gas flux measurements

Methane fluxes were measured with static chambers, spread over the alpine peatland, within the areas indicated in Fig. 1A. The locations were manually selected during each field campaign, chambers were therefore not placed at exactly the same spots during different campaigns. Both the chambers and chamber bases were constructed in-house from transparent plastic (Ikea Samla boxes). The volume of the chambers was 5 L, the volume of the bases 2 L. Bases were only used during summer campaigns and were placed in the soil minimum 1 hour before measurements. When no chamber bases were used, the chambers were weighed down with 725 ml water bottles to ensure a good contact between the chamber edge and the soil. After installation, 4 (spring and summer) or 6 (winter) discreet gas samples were collected over 30 – 60 min chamber deployments. Each gas sample was collected through a sampling port consisting of a butyl stopper with a long (10 cm) needle which’ tip was located in the central part of the chamber. The air inside the chamber was mixed prior to each sample collection. Samples (15 ml) were used to flush and then fill one 3 ml exetainer vial with 3 ml overpressure. The exetainers were then transported to the lab and CO2 and CH4 were analysed using a gas chromatograph (Thermo Fisher SCIENTIFIC TRACE 1310, Thermo Fisher Waltham, Massachusetts, USA) equipped with two pulsed discharge ionization detectors (PDD).

Gas flux rates were calculated by linear regression. For both the linear regression analysis and the creation of the boxplots, Rstudio was used (packages dplyr and ggplot2, plus readr and RColorbrewer for enhanced visualisation(Hadley Wickham, 2016; Yarberry, 2021)). Only linear regression results with an R^2^ > 0.5 (Fig. 2) or an R^2^ > 0.7 (Fig. 3) were included in the boxplots, others were discarded. In winter, emission and uptake rates were much lower than in summer, and rates closer to zero are less likely to produce linear regression results with high R^2^. To not eliminate these low rates, we chose to include data with an 0.7 < R2 > 0.5 for the seasonal analysis.

**Fig. 2.**
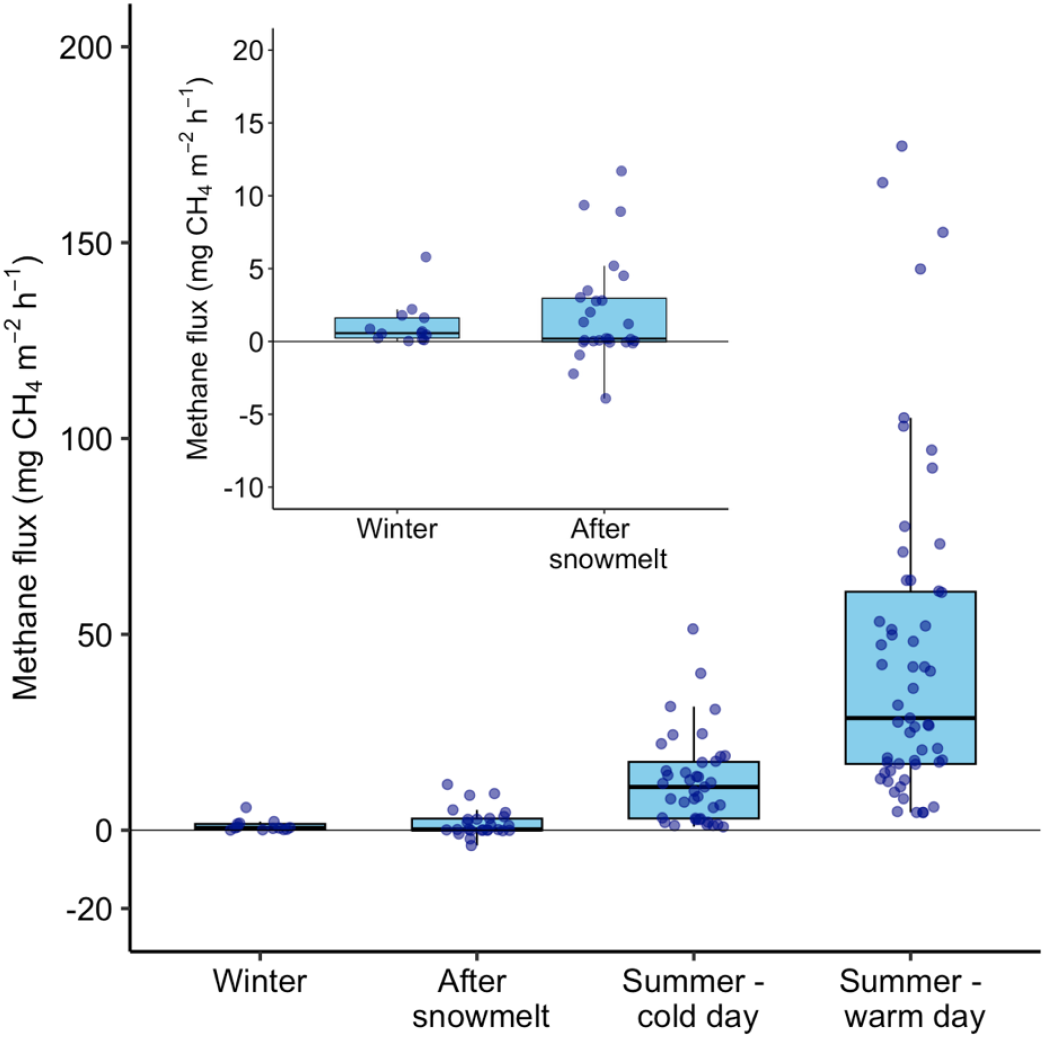
Methane fluxes from the Auenfeld peatland soil at different times of the year. The left upper corner shows the same data as in the main graph but with a different y-axis.

**Fig. 3.**
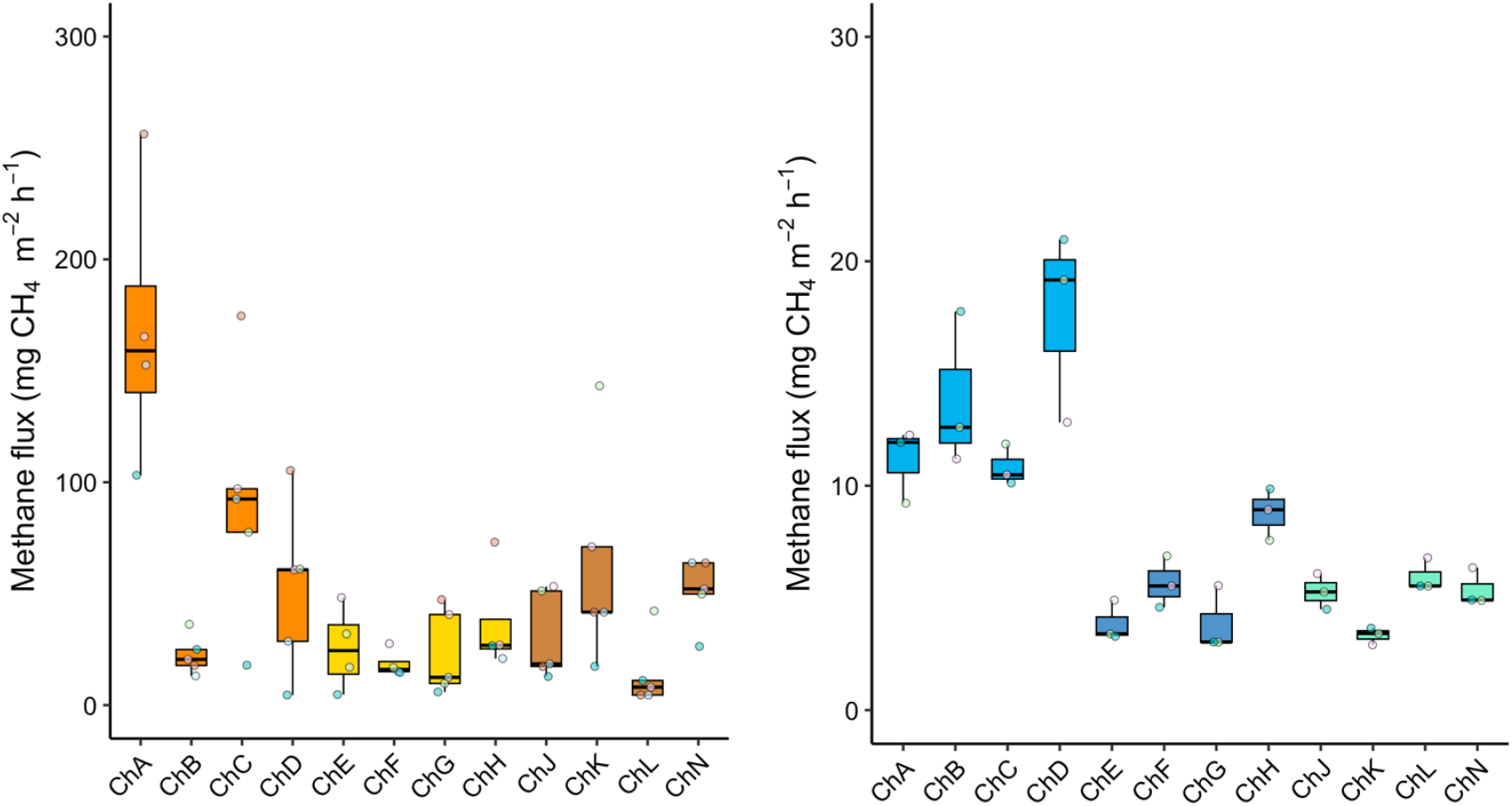
Spatial diversity of methane fluxes in the Auenfeld peatland in July 2024, during the day (left) and night (right). Note the difference in the y axis. The points are labelled according to the time of sample collection. Boxes of the same colour indicate that those chambers were located within 1 m^2^. The three m^2^-groups were 5 – 10 m apart from one another. For a picture of the setup, see Fig. A4.

The coefficient of the linear regression was corrected for chamber volume and footprint, temperature and air pressure at 1600 m or 2145 m to calculate the flux in mg CH4 m^-2^ h^-1^.

### 2.3 Soil and vegetation characteristics

The Auenfeld peatland was sampled and surveyed for soil and vegetation characteristics. The 9 locations within (and 1 location ca. 3 m outside) the peatland area that were selected for soil and vegetation analysis in August 2025 are presented in the map of Fig. 4. Each location was photographed and marked. Soil temperature was measured in situ in triplicate at 10 cm and 20 cm depth using a digital soil thermometer. Vegetation assessment was done by estimating the canopy cover of the most prominent vascular plant species and classifying them according to the Braun-Blanquet method(Westhoff and Van Der Maarel, 1978). At each site, a soil core of ca. 20 cm deep was cut from the soil with a knife. From these cores, we sampled the soil at 4 – 6 cm depth, stored it in plastic bags and kept it in the dark at 4°C until processing. In the lab, the material was dried for 24 - 36 hours at 60°C, after which it was milled using a ball-mill and used for C:N analysis on a multi N/C 2100 (Analytic Jena, Germany). Before and after drying, the samples were weighed to determine the gravimetric water content (calculated as Weightwater / Weightdry soil * 100%).

**Fig. 4.**
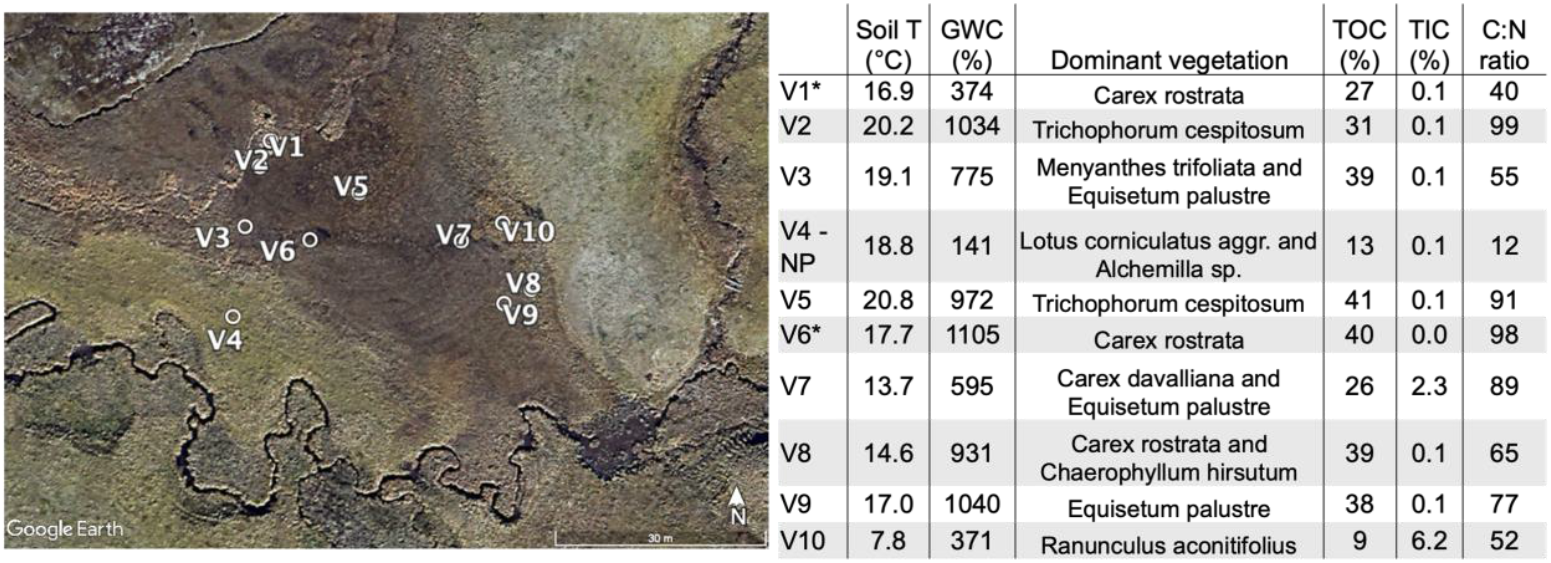
Soil characteristics across the peatland. The soil temperature reported here was measured at 10 cm depth. GWC: gravimetric water content, TOC: total organic carbon as % of dry soil weight, TIC: Total inorganic carbon as % of dry soil weight. * indicates surface water at this location. V4-NP: Non-peatland location, as reference.

### 2.4 Water sampling and analysis

Water samples in and around the peatland were collected in July 2025, after a period of 4 days of semi-continuous rain. Locations are marked in Fig. 5. The water temperature was measured at the surface with the same digital soil thermometer that was used to record soil temperatures. Water samples were collected directly from the streams into plastic syringes, filtered (0.45 μm syringe filters), placed in 12 mL exetainer vials, and stored at 4°C until analysis. Porewater sampling was performed using lysimeters (Makrorhizons, Rhizosphere Research Products, Netherlands) at 2 different depths (∼5 cm, ∼18 cm). After installing the Rhizons to the desired depths, pre-N2-flushed syringes were attached and porewater was pulled by applying a vacuum. These samples were also placed in 12 mL exetainer vials and stored at 4°C until analysis. Porewater DOC was quantified using a multi N/C 2100 (Analytik Jena, Germany) and using continuous Flow Injection Analysis (AA3, Seal analytical; UK). Major anions and cations were quantified by ion chromatography, using a ion chromatograph Compact IC Flex (Metrohm, Switzerland) equipped with an automated filtration unit and an autosampler. Certified quality controls at different concentrations were measured every 20^th^ sample and at the start and end of each run.

**Fig. 5.**
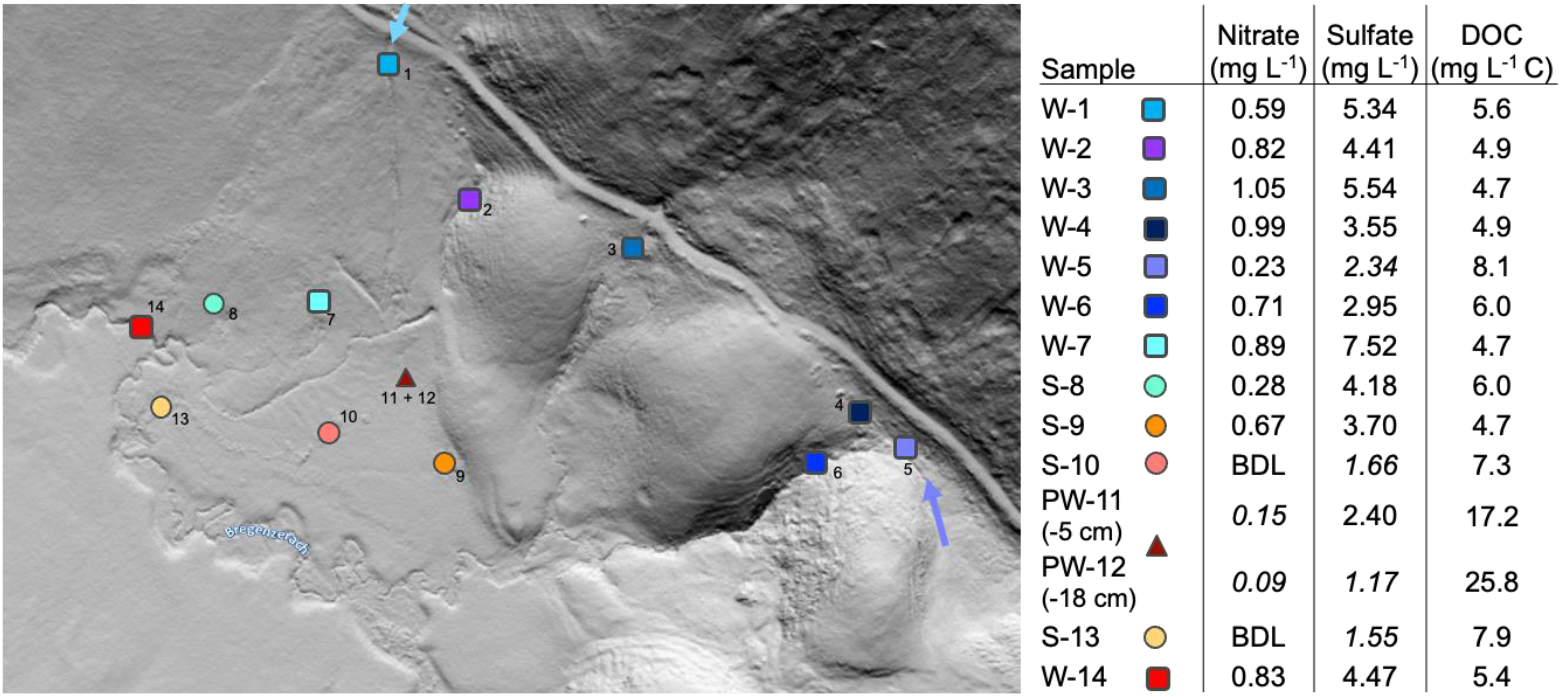
Nitrate, sulfate and dissolved organic carbon (DOC) concentrations in water samples taken in and around the Auenfeld peatland, in waterways (squares), surface water on the soil (circles), and pore water (triangles). Locations W-1 and W-5 were fed from rain-induced streams (flow origin and direction indicated by arrows). W-2, 3 and 4 were springs, meaning the water appeared from the ground at the point of sampling. BDL = below detection limit. The values in italics were outside the linear calibration range of the instrument and should therefore be interpreted with caution.

### 2.5 Microbial community analysis

One sample of the 0 – 5 cm depth layer of the peatland, collected in March 2024, was used for DNA analysis. DNA was extracted using the Powersoil DNA extraction kit (Qiagen, Germany), amplified using the general primers 515F and 806R targeting the V4 region, and used for 16S rRNA amplicon sequencing at the Tübingen Quantitative Biology Center, using their pipelines for annotation and quality control as described in (Grimm et al., 2025).

### 2.6 Statistical analyses

Besides the linear regression analyses and creation of boxplots, as described at sect. 2.2, no further statistical analyses were performed on the described data.

## 3. Results and discussion

### 3.1 Variability of summer methane emissions

The Auenfeld alpine peatland is located at 1670 m a.s.l. and is located on a slight slope (Fig. 1). It was sampled in high spatial and temporal resolution, in both a diurnal and seasonal matter. It was a strong source of methane to the atmosphere in summer (CH4 emission of 49 mg m^2^ h^-1^ on average on a warm summer day, 13 mg m^2^ h^-1^ on a cold summer day, with average temperatures and sunshine hours of the 4 days before the measurement of 15.7°C, 6.5 h and 9.7°C, 0.8 h(GeoSphere Austria), respectively), with local rates of over 150 mg m^2^ h^-1^ (Fig. 2). These local hotspots emit an amount of methane per m^2^ within one hour that other previously reported alpine peatlands emit over periods of 24 hours to one month. Other research reported summer methane emissions of 2 mg m^2^ h^-1^ (Monte Bondone, IT, 1563m)(Pullens et al., 2016); up to 8 mg m^2^ h^-1^ (Tibetan plateau, swamps and peatlands below 3000 m, reviewed by Wei et al., 2015(Wei et al., 2015) but also the Swiss Oberaar-fen, at 2320 m(Henneberger et al., 2015)); ±10 mg m^2^ h^-1^ at Goeschenen, CH (1915 m)(Liebner et al., 2012) or 3 – 30 mg m^2^ h^-1^ in a comparison study of 15 Swiss fens(Franchini et al., 2014). Our reported average rates are thus 2 – 25 times higher than in comparable systems, placing it on the high end of the peatland methane emissions spectrum. (Hirota et al., 2005) reported emissions that were on the high end for the Tibetan plateau (± 35 mg m^2^ h^-1^). Interestingly, they reported these rates after a livestock grazing experiment, which showed that grazing increased methane emissions significantly, most likely due to the resulting shortening of the vegetation. Our peatland is used for extensive grazing, unlike most of the previously investigated alpine fens and bogs. Whether grazing causes our peatland to have emission rates that are so much higher than reported in other studies needs to be determined.

Although our summer methane emissions were consistently high, there was a large spatial variation (Fig. 3). Our study design investigated both the spatial variability within 1 m^2^ and within the whole peatland. Overall, the emission rates during daytime ranged from 14 – 169 mg m^2^ h^-1^ on average (Fig. 3). Interestingly, the variation within the 1 m^2^ groups was as large as between the groups, which were 5 – 10 m apart (Fig. A4) indicating that the small-scale variation was as important as the larger scale variation between the sites. During the night, the emission rates were much lower than during the day (3.3 – 17.7 mg m^2^ h^-1^; Fig. 3) but the large spatial variation persisted, with significantly different rates between the chambers, even within the 1 m^2^ groups. The chambers that were emitting most methane during the day (Chambers A, C, K), were not the same chambers that emitted the most methane during the night (Chambers B, D).

### 3.2 Water and soil characteristics and plant community effects

#### 3.2.1 Spatial heterogeneity and methane emissions

Local differences were also observed by (Wei et al., 2015) in a swamp meadow, where they were directly correlated with soil moisture content, which ranged from 60 to 80 v%. In our peatland, the moisture content is high throughout and is therefore unlikely to be the only factor explaining the spatial heterogeneity in methane emissions, as the correlation between soil moisture and methane emissions disappears above a soil moisture content of 80%(Wei et al., 2015). Different vegetation is known to have different gas transport characteristics and may therefore influence gas dynamics locally. (Lai et al., 2014) reported different CH4 emissions based on the dominant vegetation at different sites within a Canadian peatland, similarly to (Heiskanen et al., 2021) for a Finish peatland. This effect may be further enhanced by grazing (Hirota et al., 2005). The one study on spatial variability within the European Alps focusses on the variation between different fens, not on variation within one fen(Franchini et al., 2014). We report not only high spatial variability in methane emissions, but also in soil characteristics. We sampled the soil at 4 – 6 cm depth and found a large variation in gravimetric water content (371 – 1105%) and in soil carbon and nitrogen content (TOC 9 – 41%, TIC 0.04 - 6.2%, C:N ratio 40 – 98; Fig. 4). A vegetation survey also showed large differences within the peatland, despite its small size (Fig. 4, also visible as differences in the dominant colour of the vegetation in Fig. 1C). To study the cause of this large variation on an unconventionally small scale, we measured soil temperature within the peatland and the temperature of its water sources. The soil temperature at 10 cm depth ranged from 7.8 to 20.8°C within the peatland (Fig. 4, Fig. A2), and showed a clear spatial pattern that aligned with the vegetation patterns. The water that flows upstream and downstream of the peatland has a temperature of 4.8 – 9.7°C (Fig. A2). Directly above this peatland is a steep slope (Fig. 1C) with several water springs beneath it. These springs lay just above the peatland and are present year-round. In addition to these springs, there are two streams that are only present after persistent rainfall. The springs have a constant temperature of ±5°C, the rainwater fed streams have a temperature of 8.7 – 9.7°C (Fig. A2), which may however vary over time (not measured). The temperature data suggests that the peatland is fed by the spring water, but not in a homogeneous matter. The low temperatures of the soil, despite the warm and sunny weather in the week before the measurements were taken, indicate that the spring water cools the soil in specific locations. This shows not only that certain locations receive more spring water inputs than others, but also that the residence time of the water in the peatland is most likely short. With longer residence times, the soil and soil water would be expected to warm up more. The heterogeneous pattern of water input is most likely a result of the sloping nature of the peatland and its surrounding areas.

#### 3.2.2 Solutes and soil biogeochemistry

The incoming water does not only bring colder temperatures, but also dissolved and particulate matter. Solute measurements on the spring and rainwater showed that the water contained sulfate (3.5 – 7.5 mg L^-1^ in spring water upstream of peatland). Water that had already passed through other small peatlands had lower sulfate concentrations than the spring water (2.3 mg L^-1^) and water that was located in the peatland itself as surface- or porewater had even lower concentrations (1.2 – 2.4 mg L^-1^). Nitrate concentrations were low, but followed the same pattern, with the highest concentration in the spring waters (0.8 – 1.1 mg L^-1^), a lower concentration in the water that had already been transported prior to arriving at the peatland (0.2 – 0.6 mg L^-1^) and the lowest in the water of the peatland itself (0 – 0.2 mg L^-1^). An interesting intermediate was a location in the higher part of the peatland that seems to receive high inputs of spring water (S-9, Fig. 5). It has the lowest soil temperature and the highest concentrations of both sulfate and nitrate inside the peatland, and has a much higher contribution of inorganic carbon (6.2%) than the other locations (0.1%). It is recognizable by its distinct vegetation, and the hillshade map (Fig. 1b) shows that it is the start of a small but distinguishable stream inside the peatland, therefore supporting our hypothesis that there is a heterogeneous water flow through this peatland. Observations in the field show iron precipitation on the soil surface downstream of this location. Overall, the water and soil data suggest that the spring water is an important input of redox active components (oxygen, sulfate, nitrate, possibly iron) into this peatland, but that these inputs are not equally distributed over the peatland area.

Microbial community analysis of a sample taken of the surface soil (0 - 5 cm) of the western side of the peatland shows a high relative abundance of genera involved in iron and sulfur cycling, as well as the presence of the methane oxidizing bacteria *Methylovulum psychrotolerans* and *Methylobacter* (Table S1). More microbial sampling is needed to link the microbial community to the spatial heterogeneity in methane emissions. The high relative abundance of genera involved in iron and sulfur cycling does however confirm the hypothesis that both iron and sulfur are available as redox active compounds in this peatland. Although we cannot derive the role of different electron acceptors for methane oxidation from the data collected, we expect the inputs from the spring water to play a key role in methane dynamics in the peatland, mainly by fuelling methane oxidation. More research is needed, however, to elucidate these interactions. Overall, it seems that the sloping nature of alpine peatlands plays a key role in their physicochemical characteristics and the biogeochemical cycles in the peatland area, with important implications for their greenhouse gas dynamics.

### 3.3 Snowmelt and snow season methane dynamics

The Auenfeld peatland was also sampled in May, shortly after snowmelt. During this spring period, little green vegetation biomass can be observed (Fig. A3). We measured methane fluxes in several locations within the peatland during this post-snowmelt season and again observed large spatial differences. Both methane emission and methane uptake occurred, measured fluxes ranged from emissions of 12 mg m^2^ h^-1^ to uptake rates of -4 mg m^2^ h^-1^, all within the 0.7 ha peatland area (Fig. 2). (Li et al., 2020) also reported methane uptake from a saturated alpine soil on the Tibetan plateau during the spring season, although the fluxes reported in that study were significantly lower (−0.01 to +0.04 mg CH4 m2 h) than the fluxes we report (−4 to 256 mg CH4 m^2^ h^-1^, Fig. 2). (Pullens et al., 2016) reported an increase in methane emissions after snow melt, rather than a decrease. Seasonal dynamics were also studied by (Drollinger et al., 2019) using eddy covariance tower flux data from a peatland in the lower alpine area (630m elevation). They reported positive fluxes year-round, with lowest fluxes in winter and highest in summer, with no noticeable change in spring/snow melt season.

#### 3.3.1 Snow-free patches as methane cycling locations

Previous research on methane fluxes from alpine soils, wetlands and peatlands has had a strong focus on the snow-free season. The Auenfeld peatland, although located at 1700 m and surrounded by snow-covered soils (Fig. A3), remains partially snow-free during the winter season. This phenomenon was observed in the field during our sampling campaigns (2024 and 2025, 2024 data not shown) and is also visible on historical winter aircraft images of this location (GeoAtlas Voralberg). In addition to the Auenfeld peatland site, we observed similar snow-free patches in a second peatland we studied, located at 2145 m a.s.l. (Gleirsch peatland, Tyrol, AT; Fig. A1). Soil temperature measurements showed that in both peatlands, the soil temperature at the snow-free area was consistently elevated compared to the surrounding snow-covered soil (Gleirsch: 4.6°C in snow-free peatland, 0.7°C outside the snow-free patch. Auenfeld: 2 - 4.2°C in the snow-free peatland). The temperature in the snow-free patches was close to that of the spring water that flows through the peatland and that has a constant year-round temperature of 4 - 5°C. The spring water is warming up the alpine peatland soils in winter and cooling it down in summer (Fig. A2). The snow-free areas are a result of the continuous input and relatively short residence time of the spring water in these alpine peatlands. We are the first to measure methane dynamics at these snow-free locations. At the Auenfeld peatland, the snow-free patches were sources of methane to the atmosphere (1.2 mg m^2^ h^-1^ on average, with local emissions up to 6 mg m^2^ h^-1^, Fig. 2, Fig. 6) whereas the Gleirsch wetland was a methane sink with small but consistent uptake rates (− 0.06 mg m^2^ h^-1^, Fig. 6). What causes this difference between the two studied peatlands remains unknown, and further studies are required to understand the dynamics of these snow-free patches. Previous research on snow-free peatlands has either only included locations that have an ice layer instead of snow on the soil surface (Chen et al., 2008; Gazovic et al., 2010) or were studies on artificial snow cover manipulation, which also lacked methane flux measurements(Bombonato and Gerdol, 2012; Robroek et al., 2013). Although the Auenfeld peatland methane emissions are 10-fold lower in winter than in summer, they do add up to a total winter methane release of 1670 mg m^2^ (calculated using a winter duration of 180 days, with a careful estimate of 8 hours per day of methane release, as we have not studied the nightly winter methane fluxes). The effect of the snow-free nature of these alpine peatlands in winter on their long-term soil carbon storage is currently unknown. Sites with early snowmelt have been shown to have higher organic matter decomposition rates than sites with late snow melt(Venn et al., 2021), but the higher soil temperatures seem to support plant and algae growth at both the Auenfeld and Gleirsch peatland (personal observations) which may enhance carbon uptake and storage.

**Fig. 6.**
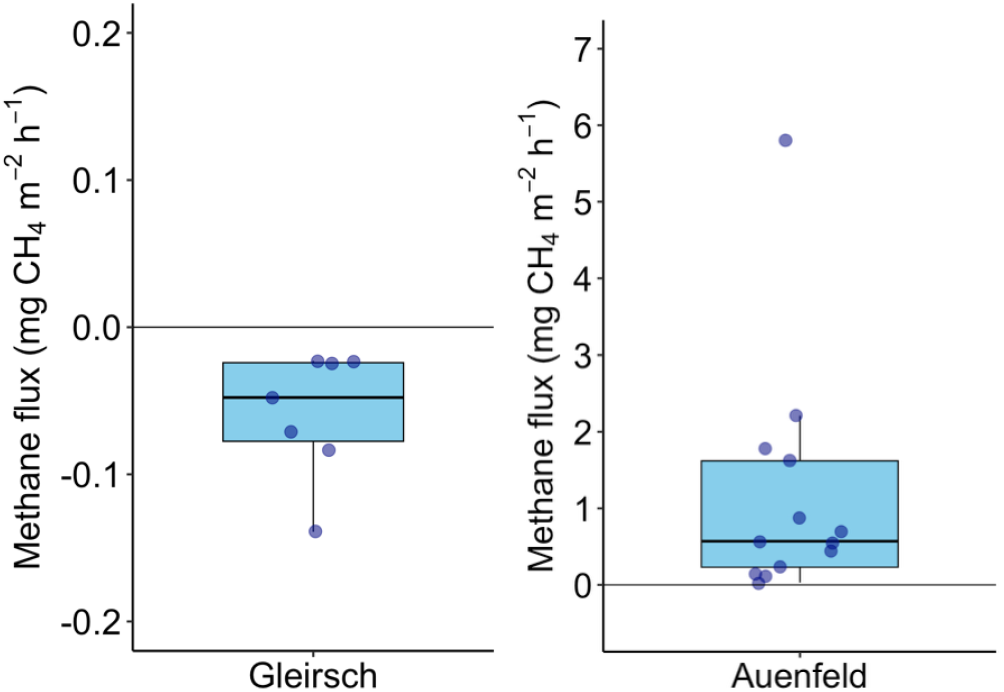
Winter methane flux in snow-free patches in alpine wetlands, in a) Gleirsch and b) Auenfeld.

Hydrological studies on hillside peatlands are scarce (Boothroyd et al., 2015; Millar et al., 2018). The relatively high spring water throughput that is a key characteristic of these systems, as shown by our research, is a result of the location and sloping nature of these alpine peatlands. In summer, the spring water inputs results in a large spatial heterogeneity in methane emission rates and in soil and vegetation conditions. In winter it results in snow-free areas that can either be methane emitters or methane sinks. The snow-free patches that we observed in the Auenfeld and Gleirsch peatlands are most likely a common occurrence in peatlands of the European Alps. Data on winter conditions in alpine peatlands is however lacking. Overall, the high variation in soil characteristics, vegetation, and methane fluxes within this small alpine peatland in both a spatial and temporal sense shows that alpine peatlands are complex and poorly understood systems that require further study.

## 4. Data availability

All data and R scripts will be made available in the public repository Zenodo upon publication.

## 5. Author contribution

SvG designed the study, wrote the first draft, and lead the data interpretation. FJ and SK performed field experiments and laboratory analyses. AK contributed to the experimental design and the writing and proofreading of the manuscript. The authors declare that they have no conflict of interest.

## 6. Acknowledgements

We would like to thank Franziska Schaedler and Lars Grimm for laboratory assistance, as well as the various members of the Geomicrobiology group who contributed to the field study. We would also like to thank the communities and municipalities of the field sites for the support.

## A Appendix

**Table A1.** Microbial community in Auenfeld alpine peatland soil (0 – 5 cm). Only species with a relative abundance of >0.7% of all 16S rRNA sequences are shown, except for the methanogens.

|  | % of 16S<br>rRNA<br>copies |  |
| --- | --- | --- |
| *Thiobacillus denitrificans* | 9.9 |  |
| *Geothrix* | 5.6 |  |
| *Rhodoferax* | 7.9 |  |
| *UBA6024 sp002429385* | 2.8 |  |
| *Flavobacterium* | 3.8 |  |
| *Sulfurimicrobium lacus* | 2.6 |  |
| *Sulfuricurvum sp028710345* | 2.1 |  |
| *Sulfuriferula plumbiphila* | 2.1 |  |
| *Magnetospirillum* | 1.8 |  |
| *Trichococcus* | 1.7 |  |
| *Methylovulum psychrotolerans* | 1.7 | Methane oxidizing bacterium |
| *UBA11358* | 1.6 |  |
| *Phenylobacterium* | 1.6 |  |
| *Pleomorphomonas* | 1.3 |  |
| *Burkholderiaceae* | 2.2 |  |
| *Methylobacter* | 1.3 | Methane oxidizing bacterium |
| *Sphingomonas* | 1.2 |  |
| *Propionivibrio soli* | 1.0 |  |
| *0-14-0-80-60-11 sp016875015* | 0.9 |  |
| *Rhizobiaceae* | 0.9 |  |
| *Limisphaerales* | 0.9 |  |
| *Clostridium_AD* | 0.7 |  |
| *Methylophilus* | 1.4 | Methylotroph |
| *Methanothrix* | 0.4 | Methanogen |
| *Methanoregula* | 0.4 | Methanogen |
| Other | 42.5 |  |

**Fig. A1.**
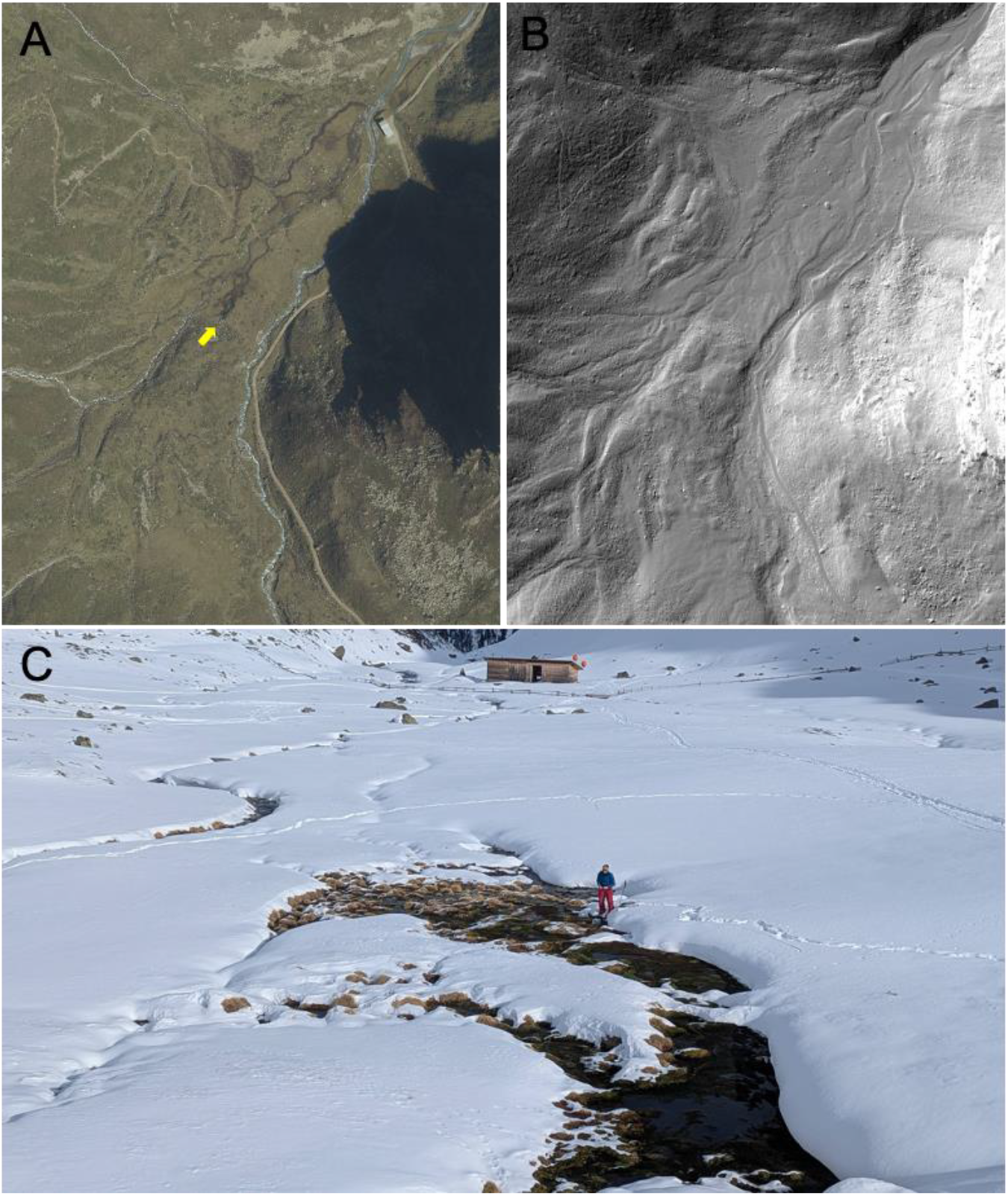
Aerial image and hillshade map of the Gleirsch alpine peatland (A and B) and the snow free patch in winter (C). The location and direction of picture C is indicated in map A with a yellow arrow. Data sources: Google Earth Pro (A) and GeoAtlas Voralberg(GeoAtlas Voralberg, n.d.) (B).

**Fig. A2.**
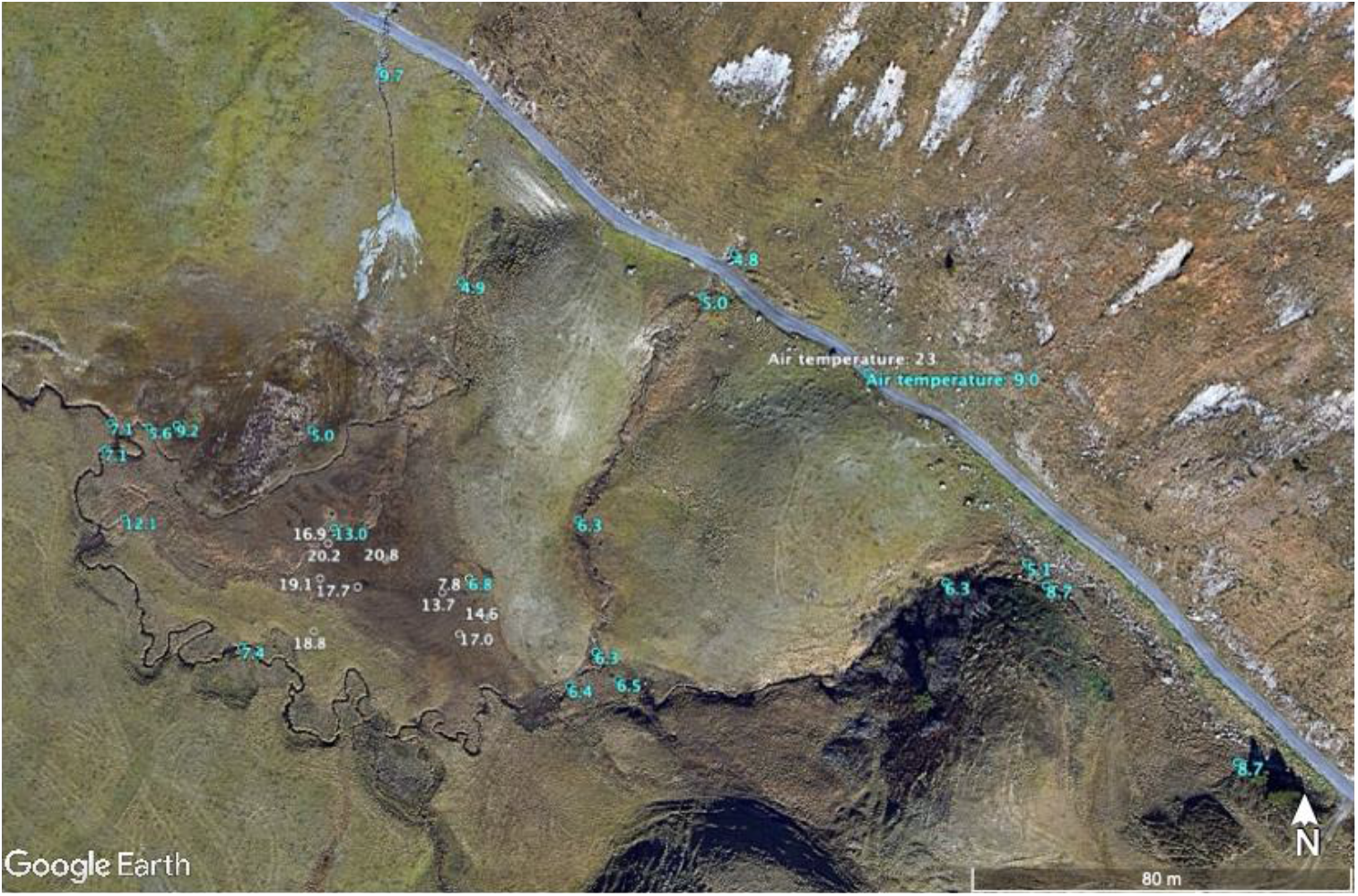
Temperature of water (in blue) and soil (in white; measurements at 10 cm depth), measured on two separate days in August 2025. The water temperature measurements were conducted after several days of rain.

**Fig. A3.**
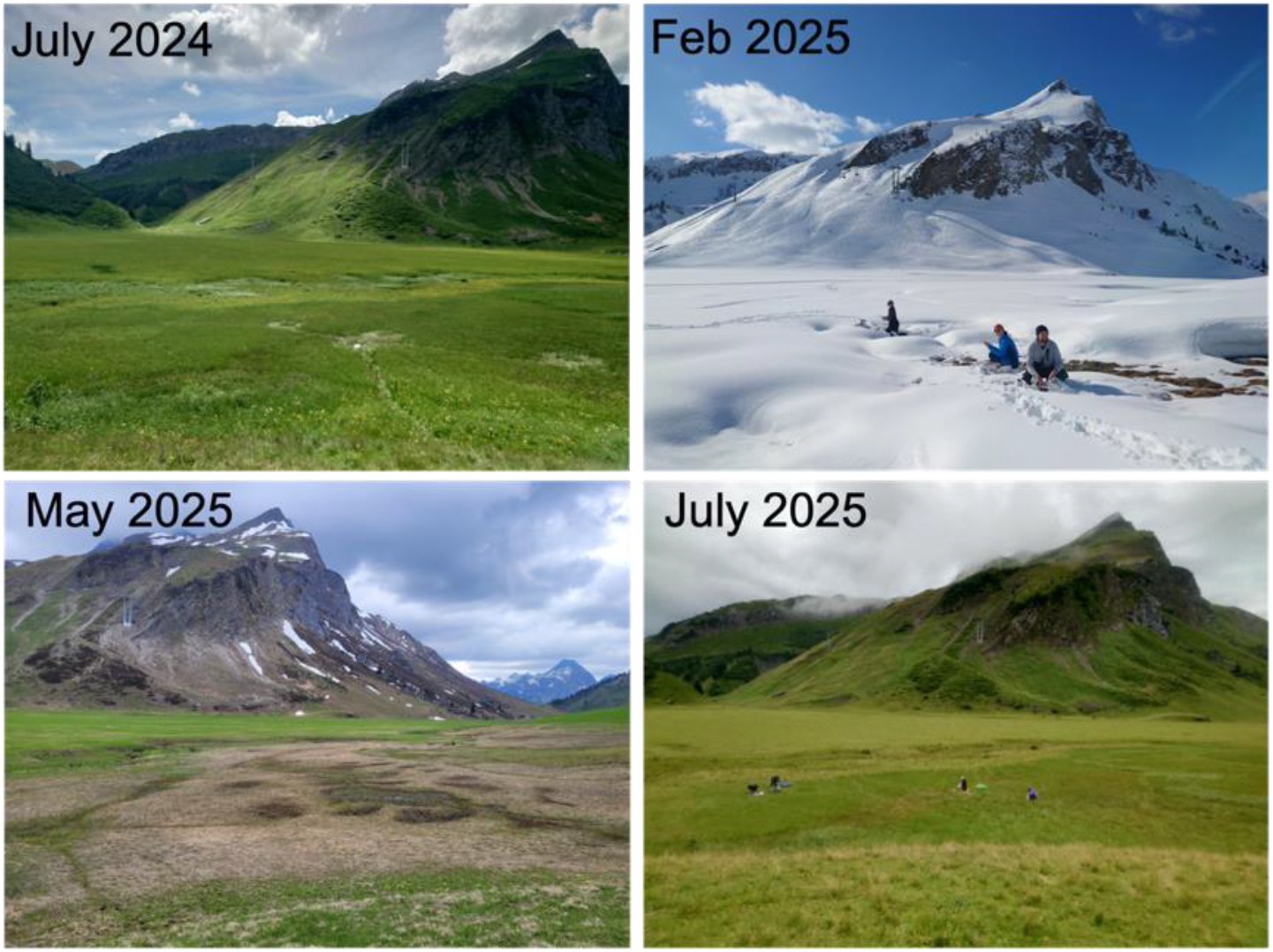
The Auenfeld alpine peatland sampling site in different seasons.

**Fig. A4.**
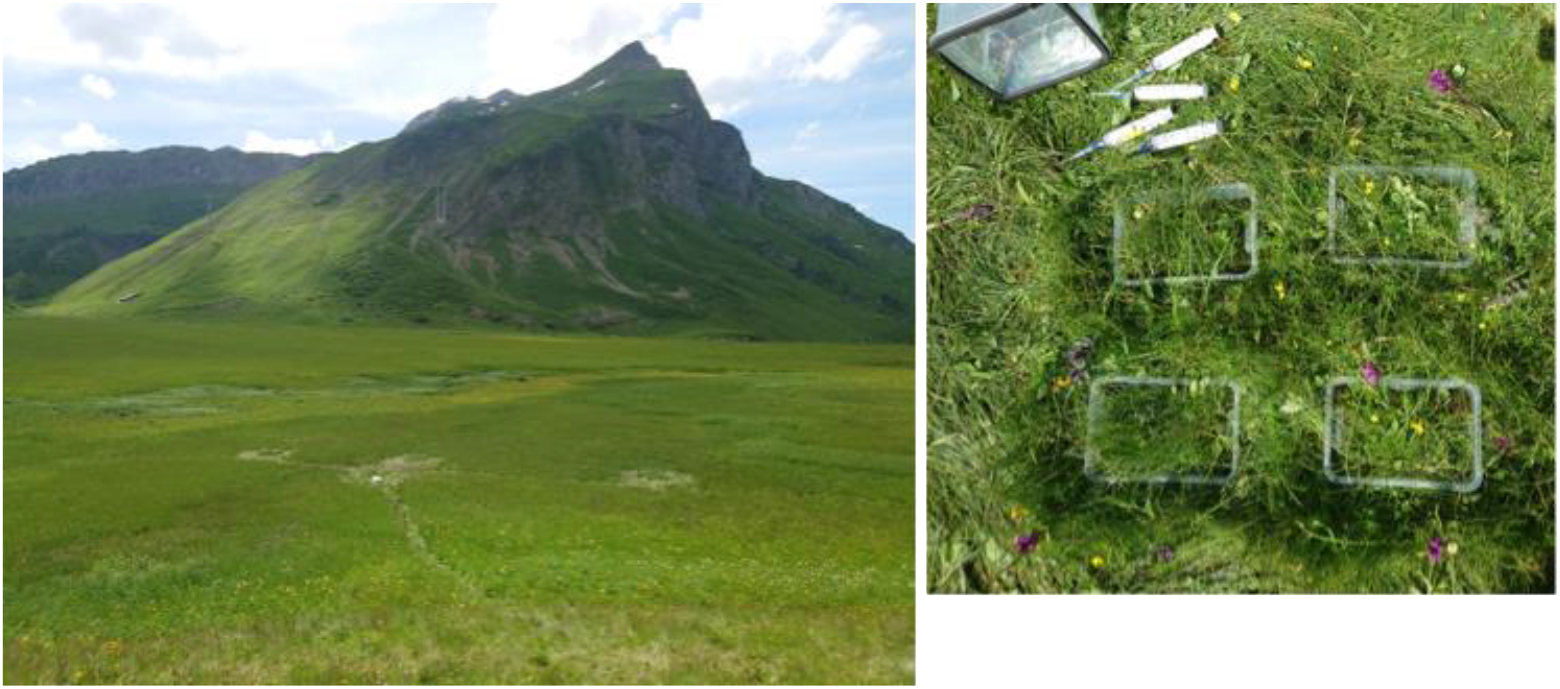
Pictures of the group-wise setup of the chambers within the Auenfeld alpine peatland. Left: overview with the three groups visible. Right: the setup of the 4 chambers within one group.

## References

Bombonato, L. and Gerdol, R.: Manipulating snow cover in an alpine bog: Effects on ecosystem respiration and nutrient content in soil and microbes, Clim Change, 114, 261–272, 10.1007/S10584-012-0405-9/TABLES/2, 2012.

Boothroyd, I. M., Worrall, F., and Allott, T. E. H.: Variations in dissolved organic carbon concentrations across peatland hillslopes, J Hydrol (Amst), 530, 372–383, 10.1016/J.JHYDROL.2015.10.002, 2015.

Chen, H., Yao, S., Wu, N., Wang, Y., Luo, P., Tian, J., Gao, Y., Sun, G., Chen, C. :, Yao, S., Wu, N., Wang, Y., Luo, P., Tian, J., Gao, Y., and Sun, G.: Determinants influencing seasonal variations of methane emissions from alpine wetlands in Zoige Plateau and their implications, J. Geophys. Res, 113, 12303, 10.1029/2006JD008072, 2008.

Chen, H., Wu, N., Wang, Y., Zhu, D., Zhu, Q., Yang, G., Gao, Y., Fang, X., Wang, X., and Peng, C.: Inter-Annual Variations of Methane Emission from an Open Fen on the Qinghai-Tibetan Plateau: A Three-Year Study, PLoS One, 8, e53878, 10.1371/JOURNAL.PONE.0053878, 2013.

Drollinger, S., Maier, A., and Glatzel, S.: Interannual and seasonal variability in carbon dioxide and methane fluxes of a pine peat bog in the Eastern Alps, Austria, Agric For Meteorol, 275, 69–78, 10.1016/J.AGRFORMET.2019.05.015, 2019.

Franchini, A. G., Erny, I., and Zeyer, J.: Spatial variability of methane emissions from Swiss alpine fens, Wetl Ecol Manag, 22, 383–397, 10.1007/S11273-014-9338-6/TABLES/5, 2014.

Gazovic, M., Kutzbach, L., Schreiber, P., Wille, C., and Wilmking, M.: Diurnal dynamics of CH4 from a boreal peatland during snowmelt, Tellus B Chem Phys Meteorol, 62, 133–139, 10.1111/J.1600-0889.2010.00455.X;WGROUP:STRING:PUBLICATION, 2010.

GeoAtlas Voralberg: Karten und Luftbilder, n.d. GeoSphere Maps: GeoSphere Austria: Messstationen Tagesdaten v2, n.d.

Grimm, H., Lorenz, J., Straub, D., Joshi, P., Shuster, J., Zarfl, C., Muehe, E. M., and Kappler, A.: Nitrous oxide is the main product during nitrate reduction by a novel lithoautotrophic iron(II)-oxidizing culture from an organic-rich paddy soil, Appl Environ Microbiol, 91, 10.1128/AEM.01262-24/SUPPL_FILE/AEM.01262-24-S0001.PDF, 2025.

Hadley Wickham: ggplot2: Elegant Graphics for Data Analysis, J R Stat Soc Ser A Stat Soc, 174, 245–246, 2016.

Harbert, B. L. and Cooper, D. J.: Environmental drivers of subalpine and alpine fen vegetation in the Southern Rocky Mountains, Colorado, USA, Plant Ecol, 218, 885– 898, 10.1007/S11258-017-0737-7/TABLES/3, 2017.

Heiskanen, L., Tuovinen, J. P., Räsänen, A., Virtanen, T., Juutinen, S., Lohila, A., Penttilä, T., Linkosalmi, M., Mikola, J., Laurila, T., and Aurela, M.: Carbon dioxide and methane exchange of a patterned subarctic fen during two contrasting growing seasons, Biogeosciences, 18, 873–896, 10.5194/BG-18-873-2021, 2021.

Henneberger, R., Cheema, S., Franchini, A. G., Zumsteg, A., and Zeyer, J.: Methane and Carbon Dioxide Fluxes from a European Alpine Fen Over the Snow-Free Period, Wetlands, 35, 1149–1163, 10.1007/S13157-015-0702-Y/TABLES/2, 2015.

Hirota, M., Tang, Y., Hu, Q., Kato, T., Hirata, S., Mo, W., Cao, G., and Mariko, S.: The potential importance of grazing to the fluxes of carbon dioxide and methane in an alpine wetland on the Qinghai-Tibetan Plateau, Atmos Environ, 39, 5255–5259, 10.1016/J.ATMOSENV.2005.05.036, 2005.

Kuhn, M., Olefeldt, D., Arndt, K. A., Bastviken, D., Bruhwiler, L., Crill, P., DelSontro, T., Fluet-Chouinard, E., Grosse, G., Hovemyr, M., Hugelius, G., MacIntyre, S., Malhotra, A., McGuire, A. D., Oh, Y., Poulter, B., Treat, C. C., Turetsky, M. R., Varner, R. K., Walter Anthony, K. M., Watts, J. D., and Zhang, Z.: Current and future methane emissions from boreal-Arctic wetlands and lakes, Nature Climate Change 2025, 1–6, 10.1038/s41558-025-02413-y, 2025.

Lai, D. Y. F., Moore, T. R., and Roulet, N. T.: Spatial and temporal variations of methane flux measured by autochambers in a temperate ombrotrophic peatland, J Geophys Res Biogeosci, 119, 864–880, 10.1002/2013JG002410;SUBPAGE:STRING:FULL, 2014.

Li, F., Yang, G., Peng, Y., Wang, G., Qin, S., Song, Y., Fang, K., Wang, J., Yu, J., Liu, L., Zhang, D., Chen, K., Zhou, G., and Yang, Y.: Warming effects on methane fluxes differ between two alpine grasslands with contrasting soil water status, Agric For Meteorol, 290, 107988, 10.1016/J.AGRFORMET.2020.107988, 2020.

Liebner, S., Schwarzenbach, S. P., and Zeyer, J.: Methane emissions from an alpine fen in central Switzerland, Biogeochemistry, 109, 287–299, 10.1007/S10533-011-9629-4/FIGURES/5, 2012.

Millar, D. J., Cooper, D. J., and Ronayne, M. J.: Groundwater dynamics in mountain peatlands with contrasting climate, vegetation, and hydrogeological setting, J Hydrol (Amst), 561, 908–917, 10.1016/J.JHYDROL.2018.04.050, 2018.

Pullens, J. W. M., Sottocornola, M., Kiely, G., Toscano, P., and Gianelle, D.: Carbon fluxes of an alpine peatland in Northern Italy, Agric For Meteorol, 220, 69–82, 10.1016/J.AGRFORMET.2016.01.012, 2016.

Robroek, B. J. M., Heijboer, A., Jassey, V. E. J., Hefting, M. M., Rouwenhorst, T. G., Buttler, A., and Bragazza, L.: Snow cover manipulation effects on microbial community structure and soil chemistry in a mountain bog, Plant Soil, 369, 151–164, 10.1007/S11104-012-1547-2/TABLES/3, 2013.

Turetsky, M. R., Kotowska, A., Bubier, J., Dise, N. B., Crill, P., Hornibrook, E. R. C., Minkkinen, K., Moore, T. R., Myers-Smith, I. H., Nykänen, H., Olefeldt, D., Rinne, J., Saarnio, S., Shurpali, N., Tuittila, E. S., Waddington, J. M., White, J. R., Wickland, K. P., and Wilmking, M.: A synthesis of methane emissions from 71 northern, temperate, and subtropical wetlands, Glob Chang Biol, 20, 2183–2197, 10.1111/GCB.12580, 2014.

Venn, S. E., Thomas, H. J. D., Venn, C. :, Thomas, H. J. D., and Peters, D. P. C.: Snowmelt timing affects short-term decomposition rates in an alpine snowbed, Ecosphere, 12, e03393, 10.1002/ECS2.3393, 2021.

Wei, D., Xu-Ri, Tarchen, T., Dai, D., Wang, Y., and Wang, Y.: Revisiting the role of CH4 emissions from alpine wetlands on the Tibetan Plateau: Evidence from two in situ measurements at 4758 and 4320m above sea level, J Geophys Res Biogeosci, 120, 1741–1750, 10.1002/2015JG002974;CTYPE:STRING:JOURNAL, 2015.

Westhoff, V. and Van Der Maarel, E.: The Braun-Blanquet Approach, Classification of Plant Communities, 287–399, 10.1007/978-94-009-9183-5_9, 1978.

Whalen, S. C. and Reeburgh, W. S.: Moisture and temperature sensitivity of CH4 oxidation in boreal soils, Soil Biol Biochem, 28, 1271–1281, 10.1016/S0038-0717(96)00139-3, 1996.

Yarberry, W.: DPLYR, CRAN Recipes, 1–58, 10.1007/978-1-4842-6876-6_1, 2021.

Zhang, P., Chu, C., Bu, X., Tong, M., Wang, H., Tian, Y., Dong, H., Zhou, D., Kappler, A., Van Cappellen, P., Waite, T. D., and Yuan, S.: Production and significance of Reactive Oxygen Species in the subsurface, 10.1016/j.earscirev.2025.105230, 1 November 2025.

